# Splicing Isoforms Associated with TGFβ-Induced Myofibroblast Activation

**DOI:** 10.1101/2025.10.06.680821

**Authors:** Opeoluwa Alli-Oke, Danny Bergeron, Fiona Kessai, Philippe Thibault, Mathieu Durand, Jean-Philippe Brosseau

## Abstract

Myofibroblast differentiation is a key process in developmental biology and involved in numerous physiopathology. The gene expression program orchestrating fibroblast to myofibroblast differentiation, as well as its recapitulation by TGFβ stimulation *in vitro*, is relatively well characterized. Intriguingly, it is known that the splicing isoform EDA+FN1 is a marker and driver of myofibroblast differentiation, but the alternative splicing landscape of myofibroblast is unknown. Here, we performed a high-throughput transcriptomic approach by RNA-Seq in a primary skin fibroblast line and uncover more than 250 splicing isoforms associated with TGFβ-induced myofibroblasts using two different bioinformatic pipelines. This splicing profile highlights a distinct layer of regulation when compared to the global gene expression profile of myofibroblasts. A 5 alternative splicing event (ASE) signature [*ACTN1-*19A/19B; *COL5A1*-64A/64B; *COL6A3* exon 4; *FLNA* exon 30 and *TPM1*-6a/6b] was further validated by ddPCR and AS-PCR and retrieved in publicly available RNA-Seq datasets describing other TGFβ-stimulated lung and skin fibroblasts. Surprisingly, TGFβ does not induce an EDA+FN1 splicing shift, although it stimulates global fibronectin expression. Thus, we conclude that the 5 ASEs signature may be used as putative universal myofibroblast markers and be of functional significance to myofibroblast formation and biology.

## INTRODUCTION

Following injury, fibroblasts follow a developmental program to differentiate into myofibroblasts. This way, they acquire enhanced contractile properties through the assembly of the characteristic stress fibers and additional extracellular matrix (ECM) secretion capacity necessary to regenerate normal tissue function [1]. One of the hallmarks of myofibroblast is the expression of alpha-smooth muscle actin (α-SMA). α-SMA overexpression not only serves to identify myofibroblasts but also drives their formation [2].

Injury-induced myofibroblast differentiation can be recapitulated *in vitro* by stimulating fibroblasts with the tumor growth factor beta (TGFβ), the master regulator of myofibroblast differentiation. The signaling cascade downstream of TGFβ activating the gene expression program driving the myofibroblasts state is incomplete. Canonical signal downstream of TGFβ receptors involved a phosphorylation cascade culminating in the translocation of a transcriptional complex binding SMAD-responsive elements, driving the expression of α-SMA, among others [3]. Some genes have been identified to drive myofibroblast activation, either by overexpressing or silencing them. An example is periostin (POSTN), a non-structural matricellular protein that has been studied to drive this developmental program [4]. In addition, the collagen family is another group of genes that has been well associated with myofibroblast activation. In fact, one of the universal markers for myofibroblast activation is the increase in the production of collagen types I and III [5].

Another major ECM protein is fibronectin (FN1). Despite the wide study of various genes linked with myofibroblast activation, only fibronectin extra domain A (EDA+FN1) has been identified as a spliced variant that drives the myofibroblast program [6]. Intriguingly, no other splicing isoforms have been associated with myofibroblast activation. Alternative splicing is a powerful process producing many protein isoforms from a single gene by selectively including or excluding certain exons of its mature RNA. Numerous examples have illustrated the functional impact of alternative splicing in the literature [7]. In this study, we hypothesized that there is more than the EDA+FN1 isoform that is associated with myofibroblast differentiation.

## RESULTS

### TGFβ-drive myofibroblast differentiation in primary skin fibroblasts

To confirm that TGFβ stimulation differentiates the primary skin fibroblast into myofibroblast; an immunofluorescence assay examining the α-SMA expression was performed. In comparison to untreated fibroblasts; formation of stress fibres was witnessed in the TGFβ-treated condition (Fig. 1A). Quantification of the α-SMA resulting fluorescence signal confirm a significant increase in the TGFβ-treated samples (Fig. 1B). Moreover, we monitored the increase in the mRNA expression of *POSTN* as an additional surrogate for myofibroblast formation by real-time quantitative PCR (qRT-PCR). As expected, we observed a significant increase in *POSTN* expression with samples stimulated with TGFβ compared with untreated samples (Fig. 1C).

**Figure 1.**
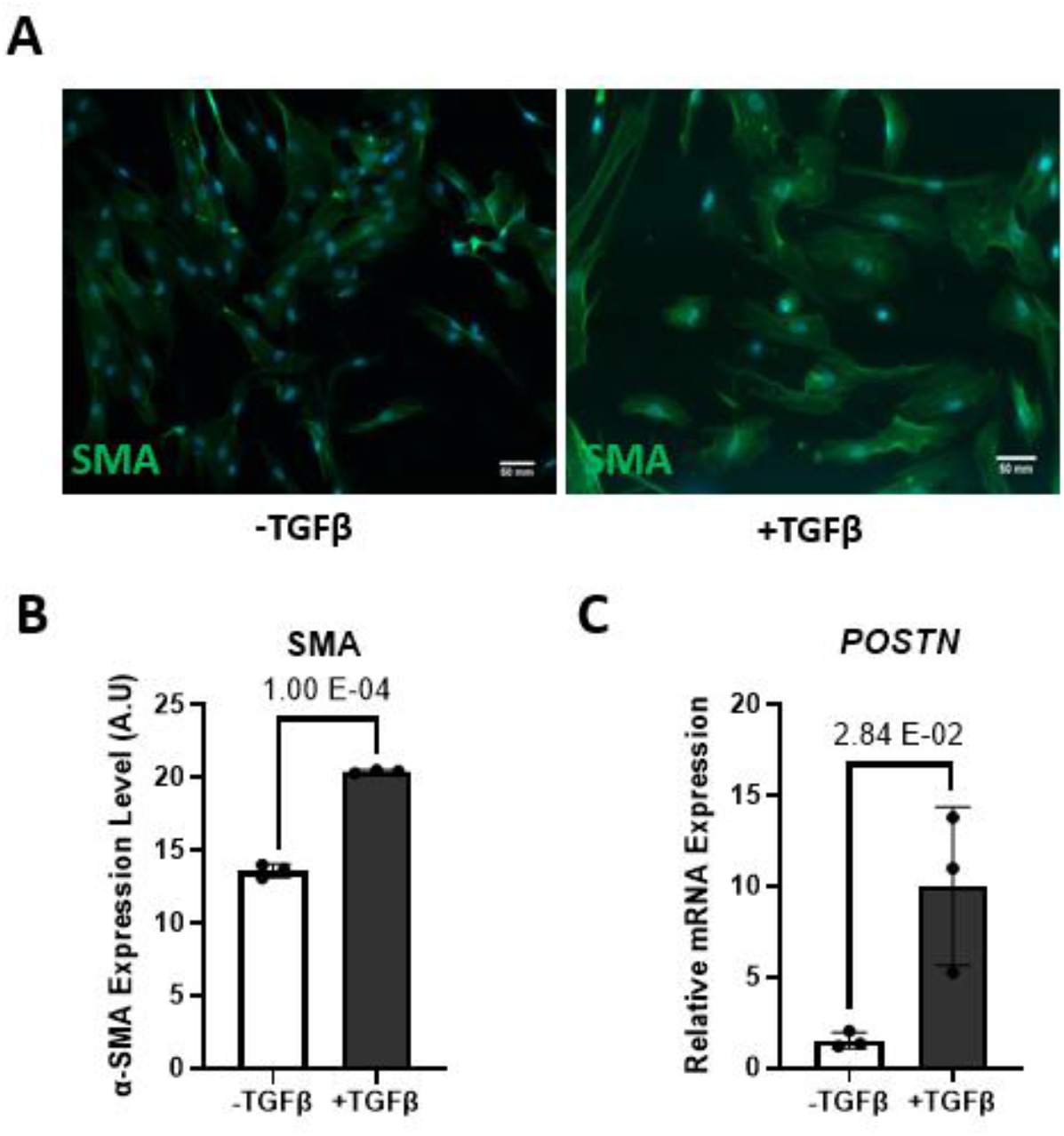
TGFβ-induced differentiation of primary skin fibroblasts to myofibroblast. **(A)** Immunofluorescence images showing α-SMA stress fibers (Green) indicative of myofibroblast after two days of TGFβ stimulation merged to DAPI (Blue). The scale bar equals 50 mm **(B)** Bar graph representing the quantification of α-SMA immunofluorescence signal from (Fig. 1A) as arbitrary units (A.U.) **(C)** Bar graph representing the relative mRNA expression of the myofibroblast marker *POSTN* by qRT-PCR.

To further confirm the myofibroblast state of the TGFβ-induced primary skin fibroblast, a genome-wide transcriptomic analysis was performed on TGFβ-treated vs untreated fibroblasts by RNA-Seq. We identified a total of 453 up-regulated genes associated with TGFβ stimulation while 1 998 genes were downregulated out of 18 910 genes (Fig. 2A-B, Table S1). Among the up-regulated genes, we noticed genes encoding collagens (e.g. *COL4A2, COL11A1, COL1A1, COL5A1, COL7A1, COL8A2*) and some involved in extracellular matrix remodelling (e.g. *MMP2, MMP8, MMP10, MMP14, LOXL2*) as expected (Fig. 2B). To determine if the 453 identified upregulated genes associated to TGFβ-induced fibroblast are significantly associate to biological processes related to myofibroblasts, we performed gene ontology analysis. As expected, biological processes associated to myofibroblast activation such as ECM-receptor interaction and focal adhesion were found (Fig. 2C). Similarly, further gene ontology analysis of the KEGG pathway also identified the ECM-receptor interaction pathways (Fig. S1A), while analysis of the molecular functions reveals the ECM structural component, fibronectin binding, and collagen binding (Fig. S1B). The cellular components analysis also consists mainly of ECM-related terms (Fig. S1C). Altogether, the phenotype and expression markers of TGFβ-induced fibroblasts are consistent with the myofibroblast state, and hence, our TGFβ stimulation procedure yields *bona fide* myofibroblasts *in vitro*.

**Figure 2.**
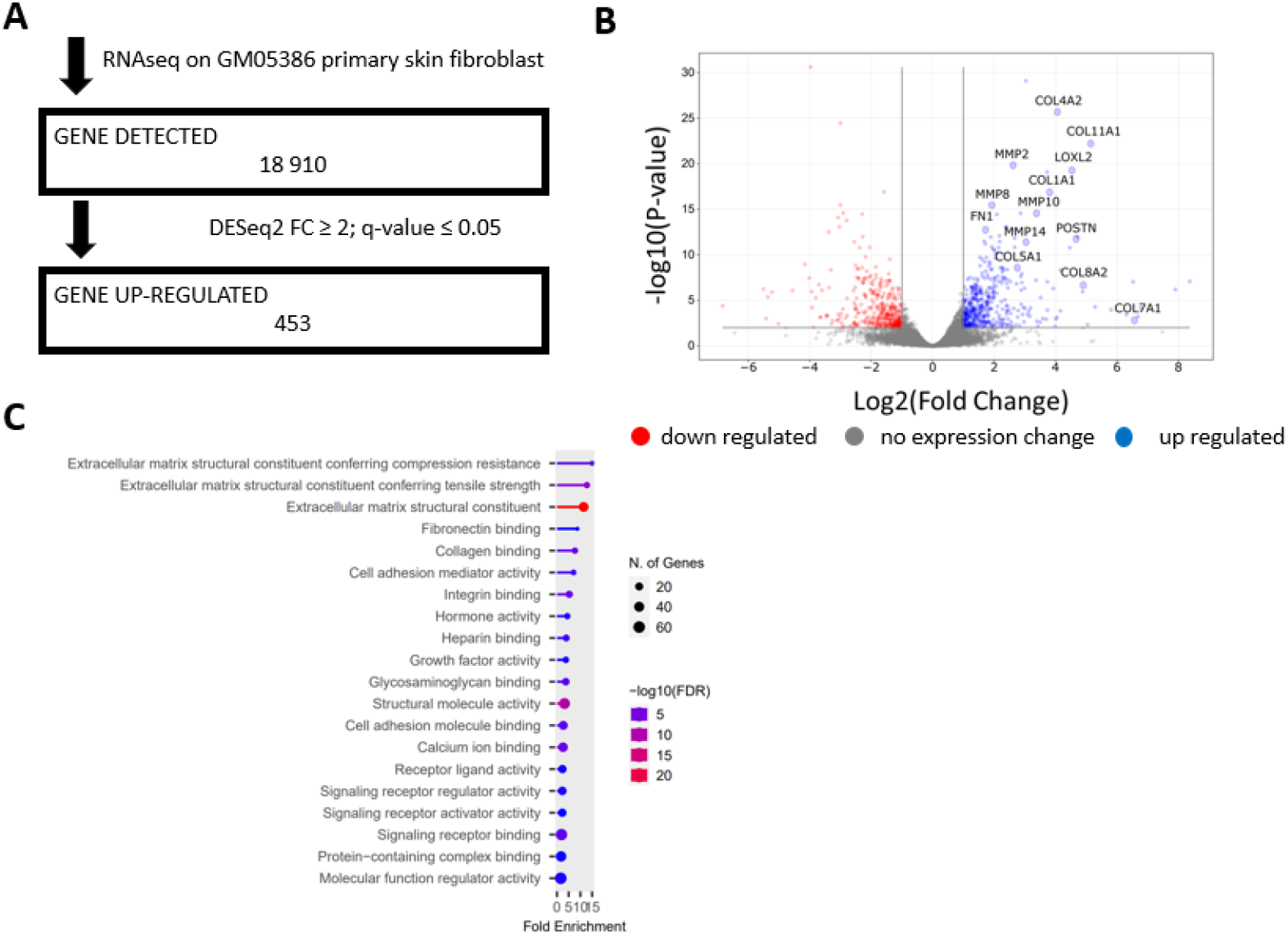
Transcriptomic analysis of TGFβ-treated primary skin fibroblast. **(A)** RNA-Seq pipeline showing the number of genes detected, cut-off parameters and TGFβ-dependent upregulated genes **(B)** Volcano plot to indicate the genes upregulated and downregulated in TGFβ-induced myofibroblast **(C)** Gene ontology for biological processes associated with TGFβ-induced myofibroblast activation.

### Transcriptome-wide splicing analysis highlights ASEs linked to TGFβ-induced myofibroblast differentiation

To identify unbiasedly any alternative splicing events (ASEs) associated with the fibroblast differentiation to myofibroblast, we re-analyzed the RNA-Seq reads we previously obtained (Fig. 2A) using two orthogonal bioinformatic pipelines (Fig. 3). The primary distinction between the two analyses is that the RNomics UdeS pipeline applies a more stringent threshold and an event-based approach [8] and rMATS provides isoform-specific analysis [9]. Using the rMATS analysis, 24 548 isoforms were detected, and 251 isoforms were found to be significantly associated with myofibroblast (Fig. 3 – left, Table S2). Out of the 30 002 ASEs detected using the RNomic UdeS pipeline, 8 ASEs were associated with myofibroblast (Fig. 3 – right, Table S3). To identify the ASEs common to both analyses, a cross-match was performed, yielding 6 ASEs that appeared in both analyses (Fig. 3, bottom). These are *ACTN1*-19A/19B; *COL5A1*-64A/64B; *COL6A3* exon 4; *FLNA* exon 30; EDA+FN1 and *TPM1*-6a/6b (Fig. 4A).

**Figure 3.**
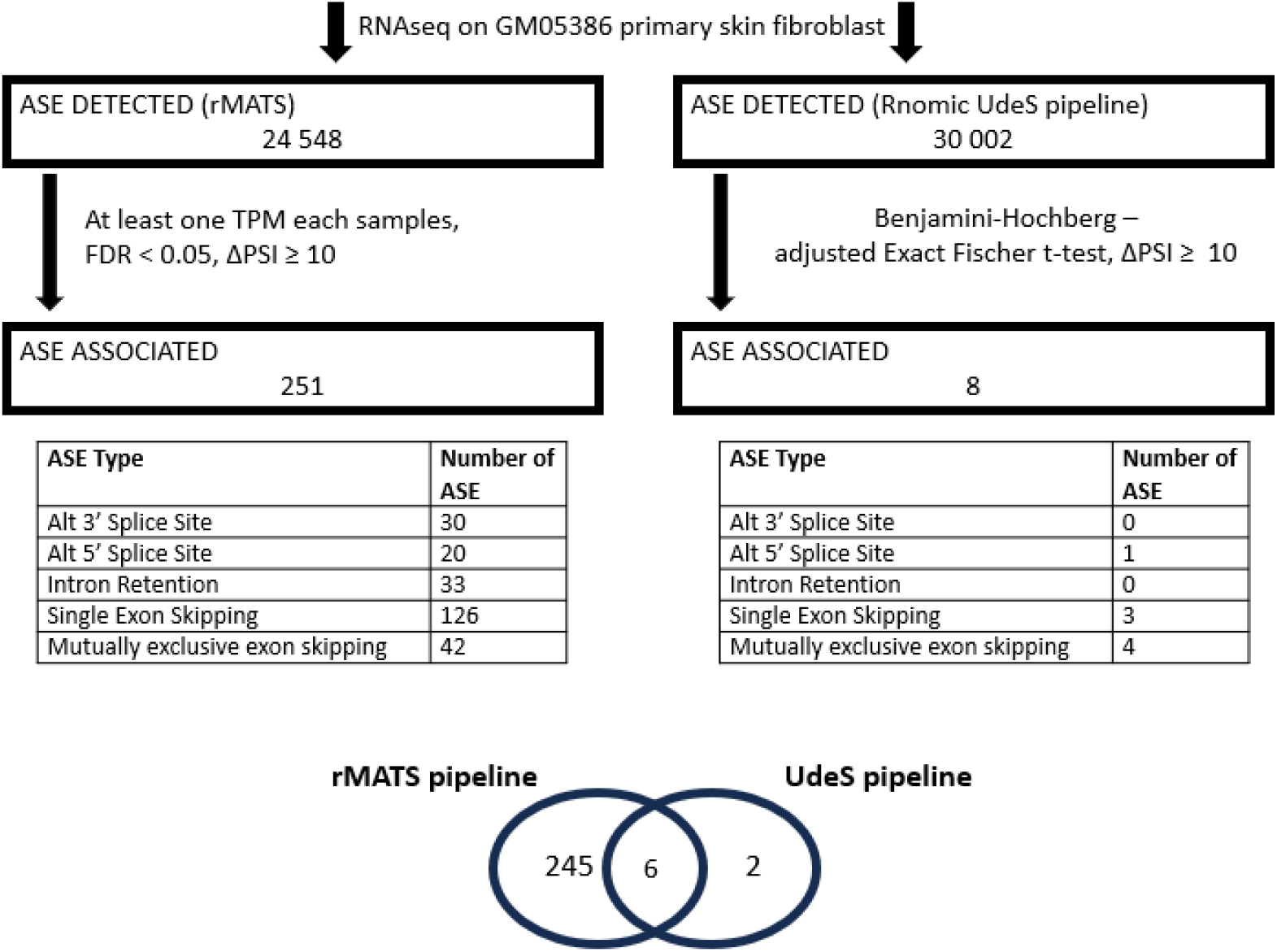
Alternative splicing isoforms associated with TGFβ-induced myofibroblast. Alternative splicing events detected by RNA-Seq and associated to myofibroblast using rMATS and our in-house pipeline. The inserted tables detailed myofibroblast-associated ASE types for each analysis and the Venn diagram at the bottom indicated the 6 common myofibroblast-associated ASEs between both analyses.

**Figure 4.**
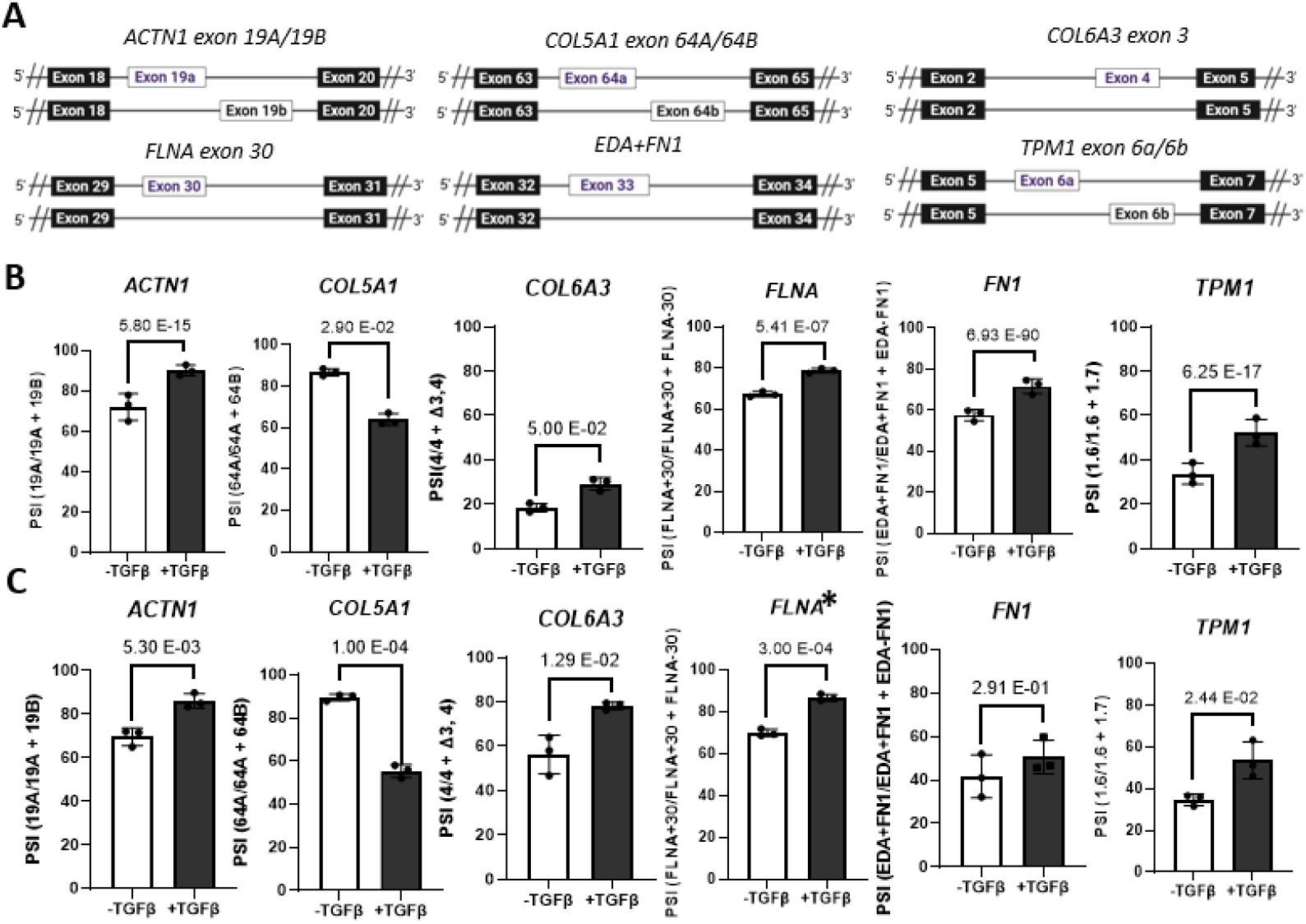
Validation of ASEs associated with TGFβ-induced myofibroblast (. **(A)** Schematic of 6 common myofibroblast-associated ASE **(B-C)** Bar graph representing the Splicing ratio of the 6 common myofibroblast-associated ASEs by (**B**) RNA-Seq and (**C**) ddPCR (* = AS-PCR).

### Experimental validation of RNA-Seq–identified ASEs associated with myofibroblast differentiation

Next, we proceeded by validating these 6 RNA-Seq identified ASEs (Fig. 4B). To do so, we performed digital droplet PCR (ddPCR) using isoform-specific primers and output the results as PSI values (Fig. 4C). AS-PCR was used for *FLNA* exon 30 because the short isoform was not detected by ddPCR. Impressively, 5 out of 6 ASEs were significantly associated with myofibroblast and importantly, the direction of splicing change was the same as initially observed by RNA-seq (Fig.4B-C). Overall, we successfully identified and validated several splicing isoforms associated with myofibroblasts.

### Evaluation of the myofibroblast gene expression signature vs the myofibroblast splicing isoform host gene signature

To determine if the global gene expression of the host gene of the 5 myofibroblast ASEs signature is differentially expressed in myofibroblast, we mined our DESeq2 analysis (Table S1) and output bar graphs specifically for the *ACTN1, COL5A1, COL6A3, FLNA*, and *TPM1* (Fig. 5A). Then, we performed ddPCR using primers targeting all spliced variants of the host genes. Overall, though a trend toward a higher expression in myofibroblast was observed, none of the gene expression could be validated by ddPCR (Fig. 5B). On a broader scale, we wanted to determine the extent of overlap between the 453 myofibroblast up-regulated genes (Table S1) and the host gene of the 251 myofibroblast ASEs (Table S2). After careful analysis, the 251 ASEs are represented by a total of 202 host genes (Table S2) and only 9 of these host genes are also up-regulated in myofibroblasts (Fig. 5C). Thus, the myofibroblast gene expression signature and the myofibroblast ASE signature is largely different.

**Figure 5.**
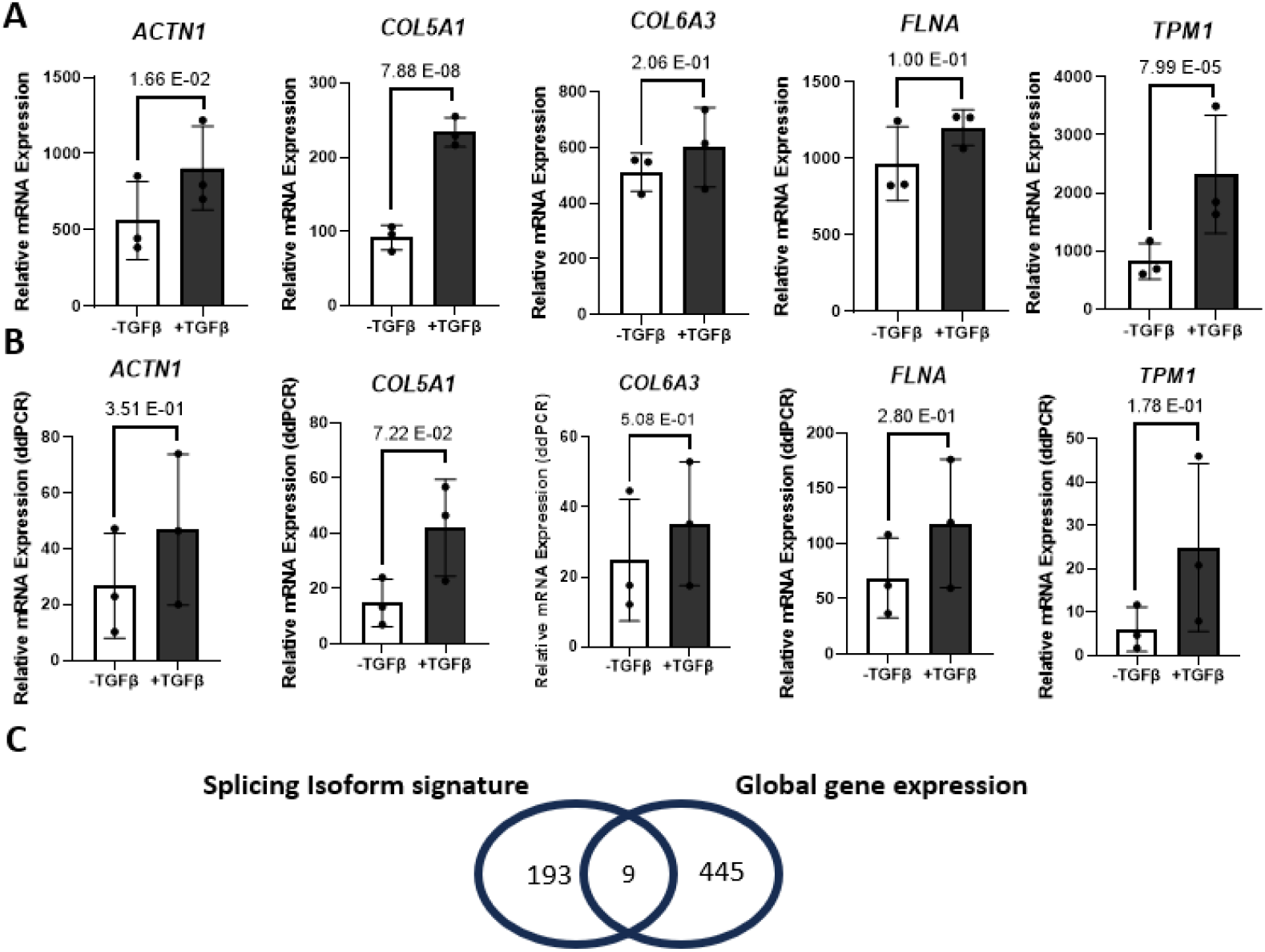
Expression level of genes with myofibroblast-associated splicing isoforms. **(A-B)** Bar graph representing the global mRNA expression level of host gene of the 5 ASEs signature by **(A)** RNA-Seq and **(B)** ddPCR data **(C)** Venn diagram to indicate common genes between 251 myofibroblast ASEs from rMATS analysis (202 host genes) and the 453 upregulated genes associated with TGFβ-induced myofibroblast activation.

**Figure 6.**
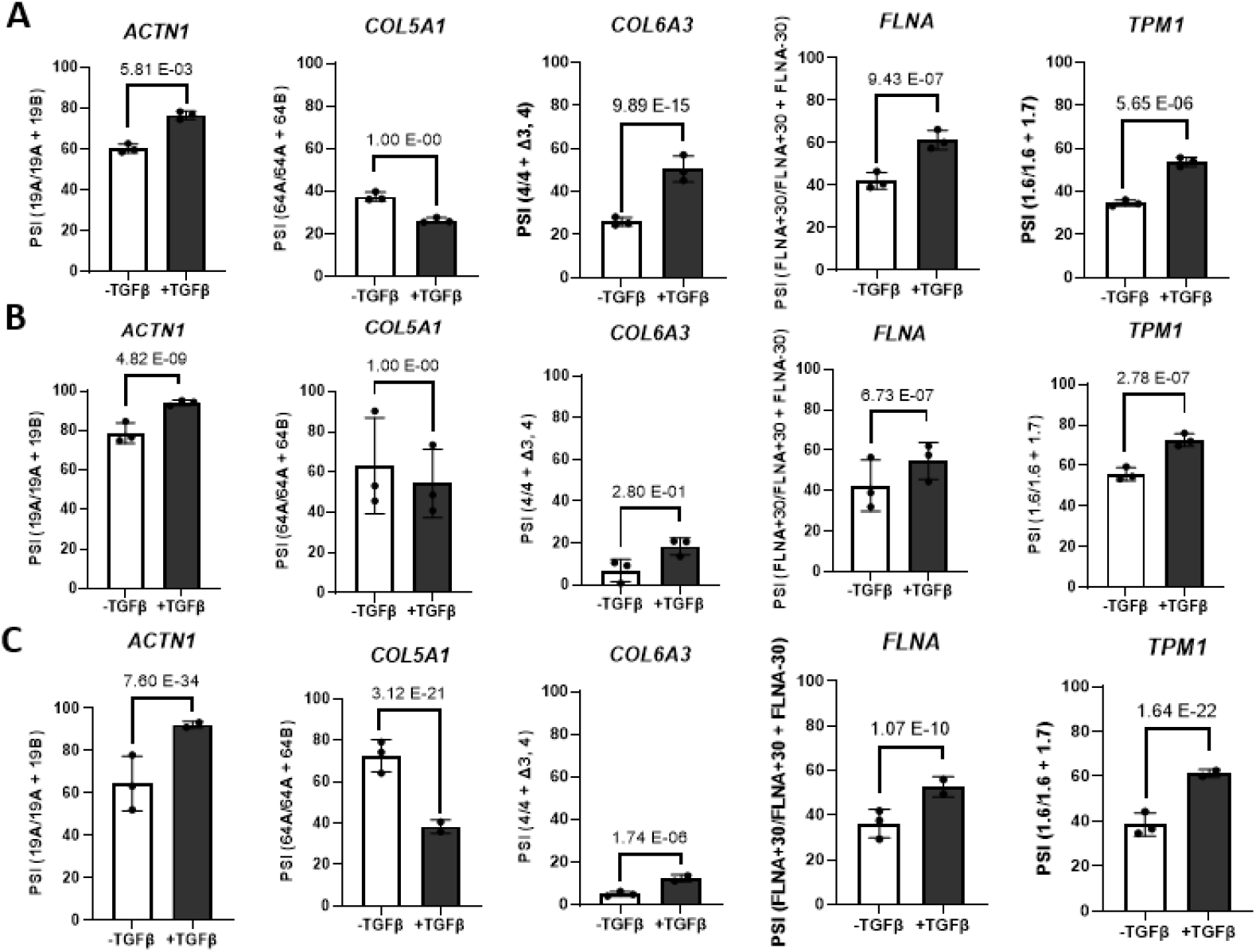
Validation of the myofibroblast-associated ASEs signature in public datasets. **(A-C)** Bar graph representing the Splicing ratio of the 5 ASE signature by RNA-Seq from the **(A)** TGFβ-induced skin fibroblasts [10], the **(B)** TGFβ-induced WI-38 lung fibroblasts from [12] and C) TGFβ-induced WI-38 lung fibroblasts from [11].

### The 5 ASEs myofibroblast signature identified is conserved across independent primary skin and lung fibroblast datasets

To confirm that the aforementioned 5 ASEs signature is not limited to the GM05386 primary skin fibroblasts we used; we searched and retrieved all possible TGFβ-induced fibroblast public RNA-Seq datasets where similar untreated controls were performed. This yields a total of three datasets: 2 using the WI-38 lung fibroblasts [10], [11] and one using skin fibroblasts [12]. Raw files were input and analyzed using our stringent in-house RNomic UdeS pipeline (Table S4), and the results were output as PSI bar graphs as previously described. Focusing on the 5 ASEs signature, and despite few outliers [splicing changes in *COL5A1* was not significant for WI-38 [10] and the skin fibroblast dataset [12], and *COL6A3* exon 4 was not significant for the skin fibroblast dataset [12]], the results indicated that the direction of change of the 5 ASEs signature is conserved across independent groups, tissues and experimental conditions. Thus, we concluded that the myofibroblast-associated splicing isoforms, especially the 5 ASEs signature, may be used as putative universal myofibroblast markers and be of functional significance to myofibroblast formation and biology.

### TGFβ-induced upregulation of fibronectin is independent of *EDA+FN1* splicing

The EDA+FN1 splicing isoform has been well studied to drive myofibroblast formation. Surprisingly, the EDA+FN1 association with myofibroblast could not be validated by ddPCR in our model system (Fig. 4C). To further investigate the EDA+FN1 spliced isoform, we designed an immunofluorescence assay experiment where we monitored the expression of EDA+FN1 using an isoform-specific antibody. As expected, the result indicates an increase in the expression of EDA+FN1 when stimulated with TGFβ (Fig. 7A). Not surprisingly, we uncover fibronectin as one of the genes up-regulated upon TGFβ stimulation in our RNA-Seq analysis (Table S1). To validate this finding, a qRT-PCR was performed, and the global expression of fibronectin was quantified (Fig. 7B). Likewise, by examining the mRNA expression level of EDA+FN1 (EDA-containing fibronectin), we observed a similar 5-fold increase between untreated samples and treated samples (Fig. 7C). However, we also find a similar value when we evaluate the expression of the EDA-FN1 (fibronectin without EDA) isoforms of fibronectin (Fig. 7D). This suggests that the TGFβ-induced fibronectin mRNA expression increase is not due to a splicing effect but to a global transcription effect. To determine the EDA+FN1 splicing index, the quantitative percent spliced in (QPSI) of the qRT-PCR was evaluated. We observed no significant change in EDA+FN1 splicing index upon TGFβ stimulation (Fig. 7E). Similar results were obtained when AS-PCR was conducted (Fig. 7F). We concluded that TGFβ does not specifically modulate EDA+FN1 alternative splicing, at least in our model system.

**Figure 7.**
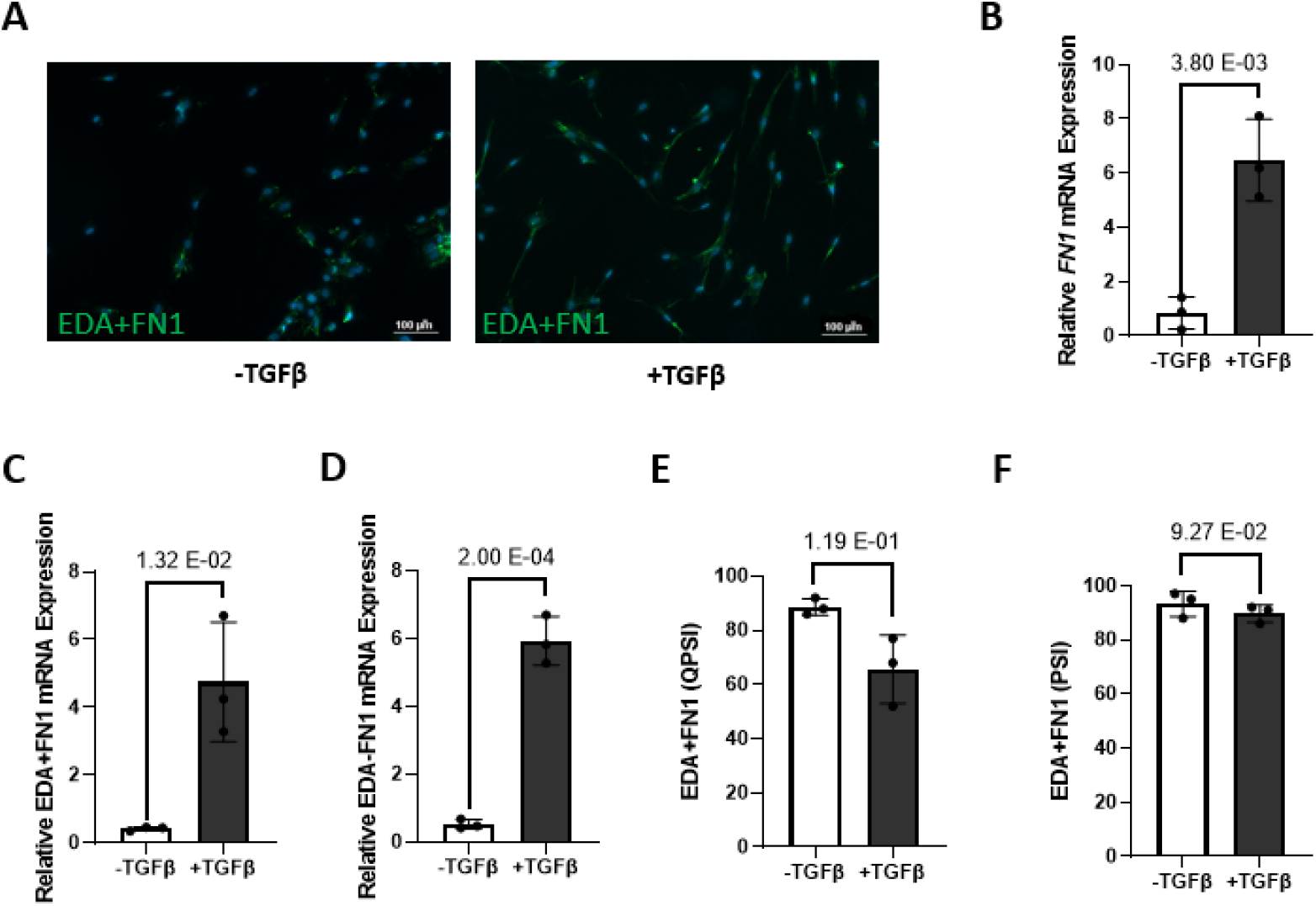
TGFβ do not specifically modulates fibronectin EDA (EDA+FN1) **(A)** Immunofluorescence images showing the expression of EDA+FN1 (Green) after two days of TGFβ stimulation merged with DAPI (Blue). The scale bar equals 100 µm. **(B)** Bar graph representing the relative *FN1* mRNA expression by qRT-PCR. **(C-D)** Bar graph representing the relative **(C)** EDA+FN1 (EDA-containing fibronectin) and **(D)** EDA-FN1 (fibronectin without EDA - right) isoform expression by qRT-PCR **(E-F)** Bar graph representing the Splicing ratio of EDA+FN1 by **(E)** qRT-PCR and **(F)** AS-PCR.

To investigate whether splicing of EDA+FN1 is impacted by TGFβ stimulation in other model systems, we interrogated the splicing analysis of TGFβ-induced fibroblasts from the public datasets analyzed previously (Table S4). The results are heterogeneous. While we observed no significant changes in the Walker dataset (Fig. S2A – upper), we noticed a PSI increase in the Liang dataset (Fig. S2A - middle) but a decrease in the Vasquez dataset (Fig. S2A, bottom). Importantly, the global fibronectin expression using (Table S5) is increased in all datasets as expected (Fig. S2B).

## DISCUSSION

Alternative splicing is an important process in gene expression, and its dysregulation has been implicated in driving many diseases and developmental programs [13,14] To investigate its role in myofibroblast formation, an RNA-Seq alternative splicing analysis was conducted in this study, and it associates more than 250 splicing isoforms. Among those, a 5 ASEs signature [*ACTN1* exons 19A/19B; *COL5A1* exons 64A/64B; *COL6A3* exon 4; *FLNA* exon 30 and *TPM1* exons 6a/6b] that was further validated by PCR emerged. We confirmed the robustness of our findings across three publicly available transcriptomic datasets. Despite differences in experimental conditions such as fibroblast origin (lung vs skin), time point (between 2 and 20 days), and TGFβ concentration (2 and 10 ng/mL), the consistency of the 5 ASEs signature across heterogeneous datasets highlights a core set of splicing changes. It supports their biological relevance to myofibroblast biology.

Unexpectedly, we could not validate EDA+FN1, a myofibroblast-associated isoform, in our model system. Unlike the 5 ASEs signature, we find EDA+FN1 to be much more variable in terms of change of splicing direction across independent groups, although we and others have found EDA+FN1 to be associated with myofibroblasts in some contexts [15], [16] and Fig. S2. Overall, it suggests that the promotion of *EDA+FN1* may be one of the mechanisms, among others, such as enhancing overall fibronectin expression.

Myofibroblasts are characterized by enhanced contractile properties through modulation of the actin cytoskeleton and enhanced ECM secretion capacity. Interestingly α-actinin 1 (*ACTN1*), filamin (*FLNA*) and tropomyosin 1 (*TPM1*) are all actin-binding proteins influencing the rigidity/stability of actin stress fibers whereas Collagen type V and type VI are myofibroblast-associated ECM proteins. However, the functional contribution of the myofibroblast-associated splicing isoforms to myofibroblast differentiation warrants further investigation. α-actinin 1 binding properties is highly regulated by calcium through its calmodulin-like domain. Calcium binding reduces conformational flexibility and impairs proper actin orientation to form a mature cytoskeleton [17]. Interestingly, the inclusion of *exon 19a* in *ACTN1* mRNA produces an α-actinin 1 isoform much more sensitive to calcium than the α-actinin 1 isoform including the exon 19b [18]. We observed an increase in inclusion frequency of exon19a over exon19b. Filamin possesses 24 immunoglobulin-like repeat units in its C-terminal, encompassing exon 30, and this domain is critical for its ability to homodimerize. Not much is known about the functional contribution of *FLNA* exon 30 to filamin activity in general, even less about its role in the context of myofibroblast in particular. Tropomyosin 1 (*TPM1*) binds and stabilizes actin filaments essential for myofibroblast differentiation. The induction of this gene increases contractility of myofibroblasts and remodelling of tissues [19]. *TPM1* is extensively alternatively spliced at exon 1, 2, 6, and 9, creating multiple isoforms. Very recently, Wu and collaborators reported that Tpm1.6, the *TPM1* variant specifically including *exon 6b*, is induced by TGFβ in renal fibroblasts [20]. Moreover, knockdown of Tpm1.6 impairs the differentiation of myofibroblasts upon TGFβ stimulation [20]. It is tempting to speculate that the 5 ASE signature may also be captured and be of significance in renal fibroblast biology. Thus, suggesting that alternative splicing adds an additional layer of regulation aiming at modulating actin filament rigidity in myofibroblasts.

Collagens are an abundant and important class of structural and functional ECM proteins secreted by fibroblasts. TGFβ stimulation leads mainly to Collagen type I expression, although myofibroblasts are also able to secrete Collagen types III, IV, V, and VI. Intriguingly, overexpression of the *COL5A1* exon 64a isoform promotes epithelial to mesenchymal transition [21].This is not what our results suggest; indicating that a switch in favor of the exon 64b isoform is associated with myofibroblast differentiation. The expression of *COL6A3* was found to be specifically secreted by cancer-associated fibroblasts [22]. Of note, *COL6A3* undergoes extensive alternative splicing of exon 3, 4 and 6. Although *COL6A3* exon 4 inclusion is associated with pancreatic cancer [23], its functional contribution has not been evaluated. Overall, several evidences suggest that the 5 ASEs signature may be of functional relevance to myofibroblast biology, although further work is needed.

## CONCLUSION

Our study reveals that alternative splicing may add an additional regulatory layer in TGFβ-induced myofibroblast differentiation, distinct from global gene expression changes. While the splicing of the canonical EDA+FN1 isoform was not modulated by TGFβ stimulation, we identified and validated 5 ASEs [*ACTN1*-19A/19B; *COL5A1*-64A/64B; *COL6A3* exon 4; *FLNA* exon 30 and *TPM1*-6a/6b] consistently associated with myofibroblast activation. These findings expand current knowledge of fibroblast biology and suggest that splicing regulation may hold functional relevance for myofibroblast biology.

## METHODS

### Cell culture and Maintenance

GM05386 primary skin fibroblasts (Coriell Institute for Medical Research) were maintained in Alpha Modification of Eagle’s medium (AMEM) supplemented with 20 mL of 10% heat activated FBS (VWR, CA76327-086), 5 mL of 5% L-glutamine (Wisent, 609-065-EL), 5 mL of 5% penicillin-streptomycin (VWR, CA45000-652), 800 uL of amphotericin B (Wisent, 450-105-QL), 5 mL of 5% sodium pyruvate (Wisent, 600-110-EL), 25 uL of 40ug/mL of fibroblast growth factor β (FGFβ) (Wisent, 511-126-QU), and 125 uL of 0.2% heparin (Sigma, H3393-50KU). The cells were passage twice a week with a seeding cell volume of 1 × 10^6^ cells. Prior to cell seeding, the T-75 flasks used for its passages are coated with 5 mL of 1ug/mL fibronectin (Sigma-Aldrich, F1141) diluted in PBS with Ca & Mg (Corning #21-030-CV) overnight at 4°C or 4 hours in 37^°^C incubator to aid adhesion and proliferation of the cells, followed by aspiration and washing of the flask with 5 mL of PBS w/o Ca & Mg (Corning #21-040-CM). For the IF assay, 1 x 10^4^ GM05386 fibroblasts were seeded in 8-well chamber slide pre-coated with poly-L-lysine (Sigma, P4707) for 30 mins at room temperature. At about 70 – 80% confluence, the cells were further processed for TGF-β Differentiation.

### TGF-β Differentiation

Primary skin fibroblast (GM05386) cells of 5 to 10 passage numbers were used for this experiment. The cells were seeded in 6-well per 1 mL of complete media (for RNA-Seq analysis), 96-well per 200 uL of complete media (for qRT-PCR analysis), and 8-well per 200 uL of complete media (for IF assay) plates and kept for 3 days until it reached 70 to 80% confluency. Afterwards, the supernatant was aspirated and fresh media without FBS was added followed by the addition of a 1 ug/mL TGF-β (Abeomics, 32-1763-10) stock to reach a final concentration of 10 ng/mL. Cells were kept in an incubator and harvested after 2 days.

### qRT-PCR

RNA extraction was performed using Qiazol reagent as per the manufacturer recommendation, and its dosage was performed using ThermoScientific Nanodrop Lite spectrophotometer. 150 ng of total RNA was converted to cDNA using the following reagents: random primer (Sigma, 11034731001), transcriptor RT reaction buffer 5X (Roche, 21-031-CV), RNase OUT, dNTPs (Wisent, 800-410), and transcriptor reverse transcriptase (Roche, 21-031-CV) and programme: 10 min @ 25°C, 30 min @ 55°C, and 5 min @ 85°C. Next, cDNA was diluted with 440 uL of RNase DNase free water and used as template for a qPCR reaction using isoform or gene specific primers and B2M as housekeeping gene (see Table S6 for primer sequences) in the 2X SyBr Green mix buffer (Quantabio, 95054-02K) under the following cycling conditions: 95°C, 3 min; [(95°C, 15sec, 60°C, 30sec, 72°C, 30sec) X 50], 72°C, 30 sec. Housekeeping normalized values were subjected to a 2^-ΔΔCt^ formula to calculate the fold changes between non-treated and treated groups.

### RNA-Seq library preparation and sequencing

RNA extraction was performed using a hybrid trizol-column protocol. Briefly, the upper aqueous phase after chloroform-trizol phase separation was then transferred and diluted with an equal volume of 70% ethanol, followed by transfer to a RNeasy mini spin column (Qiagen, 74104) as the manufacturer’s recommendation. Column-bound RNA extracts were DNAse treated using the RNAse free DNAse set (Qiagen, 79254), and their quality checked by Agilent Bioanalyzer for the RNA integrity (RIN value). Libraries were generated from 20 ng of total RNA as following: mRNA enrichment and cDNA synthesis was achieved with the NEBNext Single Cell Low Input RNA Library Prep Kit for Illumina (New England BioLabs). Adapters and PCR primers were purchased from New England BioLabs. Libraries were quantified using the KAPA Library Quantification Kits - Complete kit (Universal) (Kapa Biosystems). Average size fragment was determined using a LabChip GX (PerkinElmer) instrument. Sequencing was performed on Illumina NovaSeq by Genome Québec deposit as GEO dataset (pending).

### RNA-Seq differential gene expression - DESeq2 analysis

DESeq2 analysis was performed by the Plateforme RNomique / RNomics Plateform from the Université de Sherbrooke. Briefly, FASTQ files from this study and from GEO datasets (GSE110021, GSE252425, GSE264038) were used as input files. Reads were trimmed using Trimmomatic (V0.39, [24]) and the quality of the reads was assessed using FastQC (V0.11.9, [25]). Kallisto V0.48.0, [26] was used to align the reads to the transcriptome and to quantify the transcripts. The transcriptome of the human genome GRCh38 was created using gffread (cufflinks V2.2.1, [27]) with the Ensembl annotation and genome files (V109). Transcript abundance was combined to obtain the gene level quantification. tximport package (V1.22.0, [28] was used to summarize kallisto count estimates at the gene level. DESeq2 (V1.34) was subsequently used to identify Differentially Expressed Genes (DEGs) between the different conditions using the default Benjamini and Hochberg correction method and output as Table S1 and S5. The functional enrichment analysis was done using ShinyGO 0.77 [29], [30], [31].

### RNA-Seq Alternative splicing analysis - Rnomic UdeS pipeline

The RNomic alternative splicing analysis using the UdeS pipeline was performed by the Plateforme RNomique / RNomics Plateform from the Université de Sherbrooke as previously done in [32], with minor modifications. Briefly, FASTQ files from this study and from GEO datasets (GSE110021, GSE252425, GSE264038) were used as input files. Reads were aligned to the curated version 109 of the UCSC reference transcriptome (hg38), using the Bowtie 2 (V2.4.4) aligner [33]. The isoforms of the various genes were quantified for each condition in transcripts per million (TPM) using the RSEM tool (V1.3.3) [34]. From this transcript quantification, alternative splicing events were identified and quantified with percent-spliced-in (PSI) using the long form (L) and the short form (S) of each event. For each alternative splicing event (which may be a cassette-exon, mutually exclusive exons, alternative 5’ and 3’, etc.), a PSI value was calculated based on the ratio of the long form to the total (long + short) present in all the different isoforms containing these forms. The associated alternative splicing events were ranked using the Benjamini-Hochberg-Adjusted Exact Fischer T-test and output as Table S3 and S4.

### RNA-Seq Alternative splicing analysis - rMATS

rMATS analysis was performed by the Plateforme RNomique / RNomics Plateform from the Université de Sherbrooke. Briefly, FASTQ files from this study and from GEO datasets (GSE110021, GSE252425, GSE264038) were used as input files. Reads were trimmed using Trimmomatic (V0.39, [24]) and the quality of the reads was assessed using FastQC (V0.11.9, [25]). Trimmed reads were aligned to the human reference genome (GRCh38, Ensembl release 109) using STAR (v2.7.10a) with default parameters. Primary alignments were retained using Samtools (v1.16.1). Differential alternative splicing events between conditions were identified with rMATS (v4.1.2) using the STAR-aligned reads as input. rMATS results were filtered to retain events with a false discovery rate (FDR) < 0.05, at least one TPM in each sample, and an absolute inclusion level difference greater than 0.1 (ΔPSI ≥ 10) and output as Table S2.

### Immunofluorescence

After the TGF-β Differentiation step, supernatant was aspirated and cells were washed with 200 uL of 1X PBS w/o Ca & Mg (Corning #21-040-CM), and subsequently fixed with para-formaldehyde 4%, permeabilized with a 0.2 % Igepal CA-630 solution, and blocked with 10% donkey serum solution. Then, cells were subsequently incubated with primary antibodies [Mouse Anti-SMA (Abnova H00000059-M02), and Mouse Anti-EDA+FN (Santa Cruz, sc-59826)] and secondary antibody [Donkey anti-Mouse AF488 (Jackson ImmunoResearch, 715-545-151)]. The quantitive analysis was performed by ImageJ.

### Alternative Splicing PCR (AS-PCR) and Electrophoresis

cDNA (10 ng) was used as template for a PCR reaction using ASE specific primers (see Table S6) and the Apex Taq RED Master Mix 2X (Genesee Scientific, 42-138) under the following cycling conditions: 94°C, 3 min; 50 cycles of (94°C, 30 sec; 55°C, 30 sec; 72°C, 30 sec); 72°C, 5 min. The resulting amplicons were submitted for electrophoresis analysis using PerkinElmer LabChiP GXTouch HT. AS-PCR was performed by the Plateforme RNomique / RNomics Plateform from the Université de Sherbrooke.

### Digital Droplet PCR (ddPCR)

ddPCR was performed by the Plateforme RNomique / RNomics Plateform from the Université de Sherbrooke. Briefly, cDNA (20 ng) was used as template for a PCR reaction using isoform-specific primers or global gene primers (see Table S6) and the QX200 ddPCR EvaGreen Supermix 2X (Bio-Rad, #1864034). Each reaction mix (20 uL) was converted to droplets with the Biorad QX200 droplet generator and ran under the following cycling protocol: 95°C, 5 min; 50 cycles of (95°C, 30 sec; 59°C, 60 sec); 72°C, 5 min; 4°C, 5 min; 90°C, 5 min. Finally, resulting plate was analyzed on the Biorad QX200 reader and concentration reported as copies/uL of the final 1x ddPCR reaction using Biorad QuantaSoft software and further converted to PSI values.

### Statistical Analysis

All experimental procedures were performed with three biological replicates while analysis from public datasets are representative of at least two independent experiments. Data are represented as the mean ± SEM. RNA-Seq differential gene expression using DESeq2 was analyzed using Benjamini and Hochberg correction method, while its AS using the Rnomic UdeS pipeline was analysed using Benjamini-Hochberg-Adjusted Exact Fischer T-Test, and rMATS was analysed using a false discovery rate < 0.05. IF, qRT-PCR, and ddPCR were performed with two or three technical replicates using GraphPad software (GraphPad, La Jolla, CA, USA) and data were analyzed for significance using unpaired t-tests with the threshold of significance set at P≤0.05.

## ACKNOWLEDGEMENT

OA is an FRQ-PBEEE scholar awardee and a former recipient of the Abdenour-Nabid, MD scholarship from the Université de Sherbrooke. JPB is a recipient of the New Investigator Award from the US Department of Defense and a FRQS J2 research scholar. JPB is funded by an NSERC Discovery Grant.

## AUTHOR CONTRIBUTION

Writing: OA, JPB

Experiments: OA, FK

RNA-Seq analysis/data visualization: DB, PT

Conceptualization: JPB

Supervision: JPB, MD

**Figure S1.**
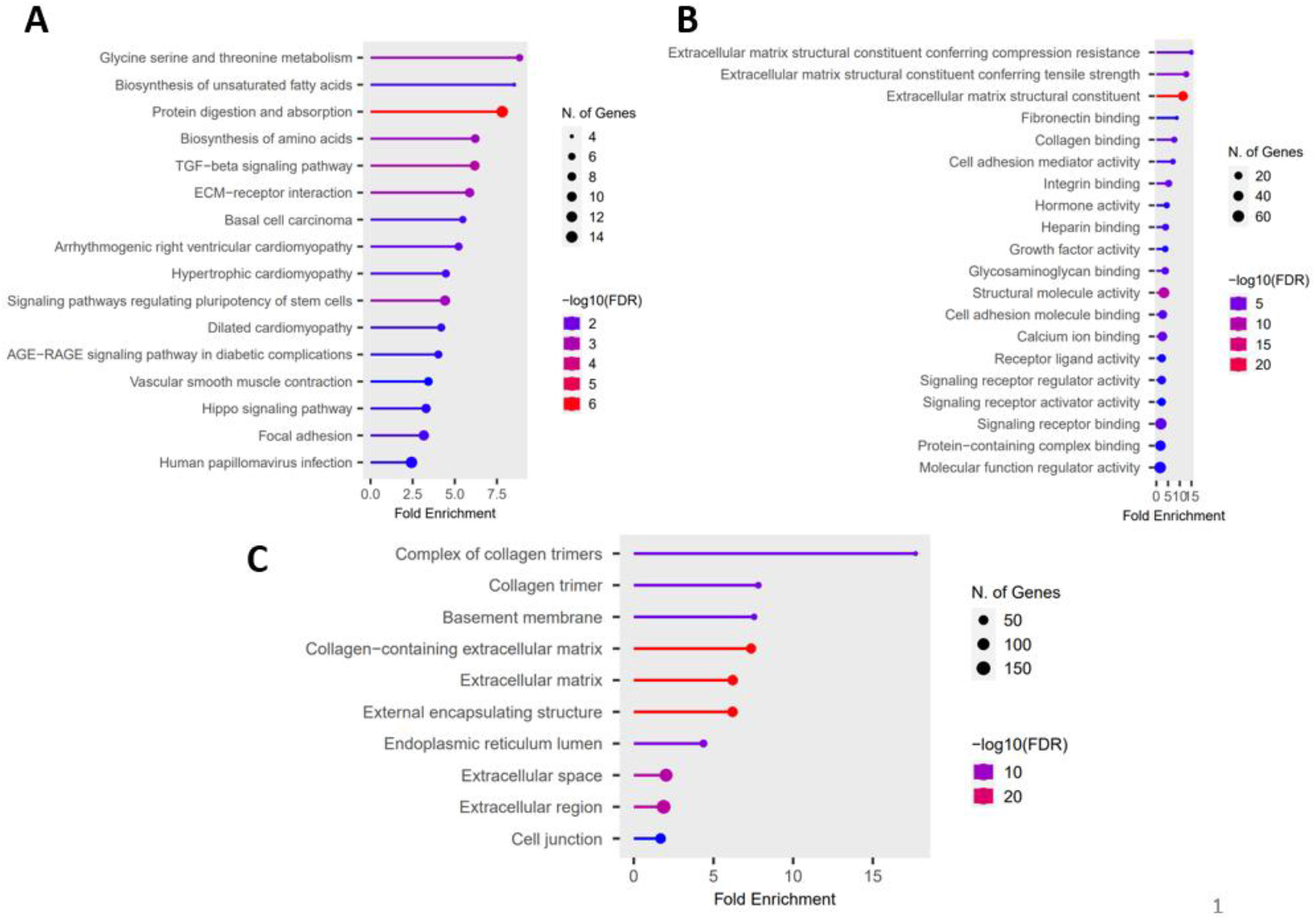
Gene Ontology analysis (up-regulated genes upon TGFβ stimulation) **(A-C)** Functional enrichment analysis of upregulated genes **(A)** Molecular functions **(B)** KEGG pathways **(C)** Cellular components.

**Figure S2. Global gene expression and Splicing ratio of fibronectin EDA**

Bar graph representing the Splicing ratio of EDA+FN1(left panels) and global gene expression of fibronectin (right panels) by RNA-Seq from the TGFβ-induced skin fibroblasts dataset [10] (upper panels), the TGFβ-induced WI-38 lung fibroblasts from [12] (middle panels) and TGFβ-induced WI-38 lung fibroblasts from [11] (bottom panels).

## SUPPLEMENTAL TABLE LEGENDS

**Table S1. DESeq2 analysis of TGFβ-induced GM053806 fibroblasts**

**Table S2. rMATS analysis of TGFβ-induced GM053806 fibroblasts**

**Table S3. UdeS Rnomic splicing analysis of TGFβ-induced GM053806 fibroblasts**

**Table S4. UdeS Rnomic splicing analysis of TGFβ-induced fibroblasts from public datasets**

**Table S5. DESeq2 analysis of TGFβ-induced fibroblasts from public datasets**

**Table S6. Primers sequence for ddPCR, qRT-PCR and AS-PCR**

